# Dynamics of organelle DNA segregation in *Arabidopsis* development and reproduction revealed with tissue-specific heteroplasmy profiling and stochastic modelling

**DOI:** 10.1101/2022.11.07.515340

**Authors:** Amanda K Broz, Daniel B Sloan, Iain G Johnston

## Abstract

Organelle DNA (oDNA) in mitochondria and plastids is vital for plant (and eukaryotic) life. Selection against damaged oDNA is mediated in part by segregation – the sorting of different oDNA types into different cells in the germline. Plants segregate oDNA very rapidly, with oDNA recombination protein MutS Homolog 1 (MSH1), a key driver of this segregation, but in contrast to mammals, we have very limited knowledge of the dynamics of this segregation within plants and between generations. Here, we combine stochastic modelling with tissue-specific heteroplasmy measurements to reveal the trajectories of oDNA segregation in *Arabidopsis thaliana* development and reproduction. We obtain and use new experimental observations of oDNA through development to confirm and refine the predictions of the theory inferred from existing measurements. Ongoing segregation proceeds gradually but continually during plant development, with a more rapid increase between inflorescence formation and the establishment of the next generation. When MSH1 is compromised, we show that the majority of observed segregation could be achieved through partitioning at cell divisions. When MSH1 is functional, mtDNA segregation is far more rapid than can be achieved through cell divisions; we show that increased oDNA gene conversion is a plausible mechanism quantitatively explaining this acceleration. We also discuss the support for different models of the plant germline provided by these observations.

## Introduction

Mitochondria and plastids are essential sites of energy transduction across eukaryotes. Originally independent organisms, they retain their own genomes (organelle DNA or oDNA; mtDNA and ptDNA respectively) encoding essential aspects of bioenergetic machinery in plants (and other eukaryotes) [Allen & Martin, 2016; Giannakis et al., 2022; Mohanta et al., 2020; Palmer et al., 2000; Clegg et al., 1994]. Plant cells typically contain populations that range from dozens to thousands of mtDNA and ptDNA molecules [Preuten et al., 2010; Greiner et al., 2020; Wang et al. 2010; Fernandes Gyorfy et al., 2021], contained within their respective organelles [MacCauley, 2013; Woloszynska, 2010; Barr et al., 2005; Johnston, 2019a]. Due to their centrality in bioenergetic, metabolic, and other cellular processes, it is essential to preserve the integrity of oDNA genes. This preservation necessitates a way of dealing with oDNA mutations and ensuring faithful inheritance of oDNA between generations.

Mutations in oDNA can give rise to heteroplasmy – a mixture of several oDNA types within a cell [Wallace & Chalkia, 2013; Stewart & Chinnery, 2015]. Across eukaryotes, developmental and genetic processes exist to limit the inheritance of heteroplasmy [Edwards et al., 2021]. In several animals, mtDNA inheritance is shaped by the so-called developmental bottleneck [Johnston, 2019b; Stewart & Chinnery, 2015; Zhang et al., 2018]. Here, cell-to-cell variance in heteroplasmy is increased in the female germline, so that individual gametes have a wide range of heteroplasmy levels. Through this increase in variance – called segregation or “sorting out” – it is then possible for some gametes to inherit lower levels of damaging mutations than the mother’s average. If gametes with high levels of such mutations are removed by selection, the mutational burden passed to the next generation is limited.

Previous work has characterised inheritance and vegetative sorting of heteroplasmy in carrot [Mandel et al., 2020]. Here, little evidence was described for segregation during plant development, with most observations involving a loss of heteroplasmy between generations. The heteroplasmy levels involved in this study were typically extreme (around 1% frequency of the minor allele), meaning that such segregation would be very hard to detect; and one notable instance was recorded of a 31% heteroplasmic offspring arising from a <1% heteroplasmic mother and father, suggesting that a mechanism for substantial amplification of minor alleles may be present. The loss of heteroplasmy upon inheritance agrees with results in Silene [Bentley et al., 2010], where only 17% of offspring retained heteroplasmy that was present in their mother.

How plants limit the inheritance of these damaging mutations is less well understood [MacCauley, 2013; Woloszynska, 2010; Barr et al., 2005; Galtier, 2011]. Although the observation of within-plant segregation of oDNA-linked phenotypes dates back over a century (and led to the discovery of cytoplasmic inheritance) [Hagemann, 2010; Greiner 2012], the quantitative dynamics and mechanisms of this segregation remain unclear. Recent experimental evidence has shown that sorting out of plant mtDNA and ptDNA is extremely rapid compared to animals [Broz et al., 2022]. This work showed that this sorting depends on *MSH1*, a gene responsible for controlling recombination activity in organelle DNA [Abdelnoor et al., 2003]. Although the precise nature and mechanism of this control is yet to be determined [Arrieta-Montiel et al., 2009; Virdi et al., 2015; Christensen, 2014], *MSH1* is required to maintain a low mutational burden in plant oDNA [Wu et al., 2020], accelerates oDNA segregation [Broz et al., 2022], and supports oDNA gene conversion [Gualberto et al., 2014; Edwards et al., 2021]. Other recombination factors including members of the *RECA* gene family also contribute to oDNA maintenance [Rowan et al., 2010; Maréchal & Brisson, 2010; Day & Madesis, 2007; Shedge et al., 2007; Miller-Messmer et al., 2012]. Theoretical work has explored the role of recombination processes in shaping plant oDNA [Atlan & Couvet, 1993; Albert et al., 1996], suggesting that gene conversion provides a strategy for oDNA segregation [Lonsdale et al., 1988; Khakhlova & Bock, 2006], with stochastic modelling showing that such segregation can occur without requiring a reduction in cellular oDNA copy number [Edwards et al., 2021]. This feature is potentially useful for plants, where, due to developmental dynamics, a germline cannot readily be sequestered and manipulated to impose a physical bottleneck. oDNA copy number in plant meristems is lower than in many animal cases [Edwards et al., 2021; Preuten et al. 2010; Wang et al. 2010; Greiner et al., 2020], but this reduction alone cannot account for the extent of segregation observed [Broz et al., 2022]. The developmental history of the plant germline differs dramatically from the animal case [Lanfear, 2018; Burian et al., 2016], and any understanding of how oDNA segregation proceeds during development necessitates an analysis approach that can both account for the developmental history underlying samples [Wilton et al., 2018; Stadler et al., 2021] and the uncertainty over different models of plant germline development [Lanfear, 2018; Kirk et al., 2013].

Here, we attempt to illuminate the dynamics and mechanisms by which plants perform this rapid sorting of oDNA heteroplasmy. We combine existing heteroplasmy measurements within and across plant generations with a stochastic phylodynamic model for cellular oDNA dynamics during plant development. We use Bayesian inference and model selection to reveal when and where cell-to-cell variability is generated; model selection and mathematical analysis reveals the likely physical mechanisms responsible for this segregation. We confirm the predictions of this model with new experimental observations, characterising the segregation dynamics of mtDNA and ptDNA within plants in unprecedented quantitative detail.

## Results

### Developmental models for heteroplasmy within and across plant generations

To use heteroplasmy measurements through developmental history to infer the dynamics of oDNA segregation, we require a quantitative model connecting the behaviour of heteroplasmy at the different developmental and generational timepoints we observe [Wilton et al., 2018; Johnston et al., 2015; Burgstaller et al., 2018; Burian et al., 2016]. We analyzed bulk tissue samples, so cell-to-cell variability cannot be directly quantified; instead, we assume that the heteroplasmy mean in a tissue sample reflects the heteroplasmy of the single cell that was the developmental ancestor of the tissue [Burian et al., 2016; Furner & Pumfrey, 1992; Irish & Sussex, 1992]. This assumption allows for any amount of segregation to occur during the development of the tissue from the precursor cell but assumes there is no systematic shift due to selection for one oDNA type over another (compatible with evidence in this system [Broz et al., 2022] and others [Mandel et al., 2020]).

Given this picture, bulk heteroplasmy samples from different tissues are interpretable as readouts of single-cell heteroplasmy in the population of stem cell precursors to each tissue. For example, mean heteroplasmy samples from three leaves are interpreted as three single cell heteroplasmy values from the (earlier) population of stem cells that gave rise to those leaves. We can then construct a developmental model inspired by the “ontogenetic phylogeny” picture tracking the relationships between cells at different developmental stages [Wilton et al., 2018]. Here, the developmental history of a set of cells is accounted for by a “cell pedigree” or “lineage tree” [Stadler et al., 2021] describing the relationship between ancestral and descended cells. Wilton et al. [2018] used such a picture to infer rates of segregation and mutation through human development given cellular profiles of the presence of different heteroplasmic variants. We will follow this philosophy but instead work with plant development and the continuous heteroplasmy level as it varies through development. This model describes and links the distributions of heteroplasmy in the estimated stem cell populations through and between generations (Fig 1A-B; see Methods). We consider three different models, corresponding to no sequestered germline, separate germline and soma developmental lineages, and a separate developmental lineage for every tissue we consider [Lanfear, 2018] (Fig. 1A). Importantly, our model considers individual heteroplasmy measurements, not their summary statistics – the sample variance, for example, would in most of our cases have a high uncertainty due to the limited number of observations in each sample. Instead of fitting a model using coarse-grained, uncertain summary statistics, we use the full set of individual heteroplasmy measurements in a parametric framework to obtain high statistical power [Giannakis et al., 2023].

**Figure 1.**
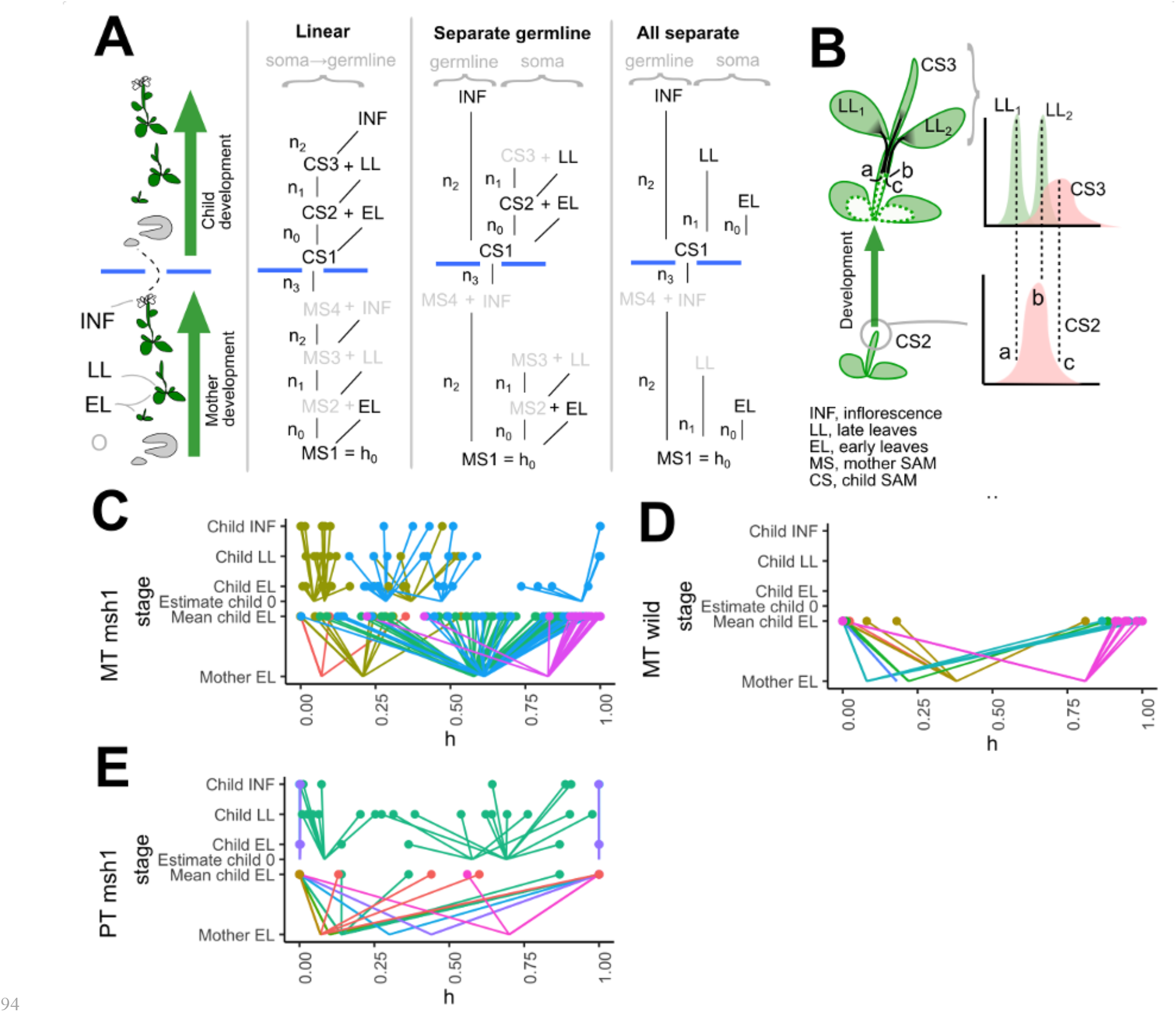
Models and data for heteroplasmy segregation in plant development. (A) Developmental models for heteroplasmy observations. MSi and CSi are the unobserved (latent) ancestral cells at different developmental stages in Mother and Child shoot apical meristem (SAM). The blue horizontal bars denote the generation of sex cells and establishment of a new generation. Greyed-out elements are unidentifiable given our observations and play no role in our model. n_i_ correspond to the number of effective segregation events (model cell divisions) at each developmental stage. (B) Example of heteroplasmy model within the linear developmental model in (A). The SAM at the CS2 stage includes cell with a distribution of heteroplasmy levels. In this example, three cells a, b, and c from this distribution, with different heteroplasmy levels, go on to be the ancestors of two late leaves (LL_1_ and LL_2_) and part of the future SAM at stage CS3. Segregation increases heteroplasmy variance as the descendants of a, b, and c develop, leading to new distributions. These may be sampled (the mean of LL_1_ and LL_2_ are recorded) or unseen (the CS3 distribution plays a latent role in our model). (C-E) Observed heteroplasmy data through development in different heteroplasmic plant families from Broz et al. [2022]: (C) mtDNA in mutant *msh1* background; (D) mtDNA in wildtype background; (E) ptDNA in mutant *msh1* background. Between-generation (upper) and within-plant (lower) observations are shown. The “fanning out” of individual sample heteroplasmies over time corresponds to increasing sample-to-sample variance.

The amount of segregation occurring between each developmental period is quantified in our model as “effective segregation events”. This is the number *n* of binomial cell divisions (and associated oDNA reamplifications) that would generate the observed heteroplasmy variance, with an effective population size *N_e_*. We use this variable rather than a “bottleneck size” or “drift parameter” [Johnston, 2019b; Wonnapinij et al., 2008] because (a) it corresponds to a biological “null model” where variance is generated by cell divisions alone (see below); and (b) because it is a convenient additive quantity, so that the effective number of segregation events describing *n_1_* events followed by *n_2_* events is simply *n_1_*+*n_2_*. We assume, based on biological observations in the *Arabidopsis* germline (see Methods), that *N_e_*= 50 for mtDNA [Wang et al., 2010; Preuten et al., 2010] and 7 for ptDNA (the latter corresponding to 7 genetically homogeneous organelles [Greiner et al., 2020; Scarcelli et al., 2016]). We adopt binomial cell divisions and reamplification as a convenient null model with some empirical support [Johnston et al., 2012; Johnston et al., 2015], although mtDNA partitioning in yeast has been observed to be controlled to a tighter extent [Jajoo et al., 2016].

To learn the likely mechanisms of oDNA segregation in real plants, we begin with the dataset from Broz et al. [2022], labelled by different developmental stages (Fig. 1C-E). These stages are early-emerging leaves (EL, fully expanded between 4-6 weeks of growth), late-emerging leaves (LL, upper rosette leaves that were fully expanded after 8 weeks of growth), and inflorescences (INF) (Fig. 1A; see Methods), reflecting tissues generated progressively later in development from the SAM. These data include observations of mtDNA heteroplasmy in wild type and *msh1* mutant backgrounds, and ptDNA heteroplasmy in the *msh1* mutant. All wild type lines measured were homoplasmic in ptDNA, likely due to the high rate of wild type segregation [Broz et al. 2022].

### Generation of heteroplasmy variance across tissues and between generations

We first aim to infer the number of effective segregation events at each developmental stage in Fig. 1. We used reversible jump Markov chain Monte Carlo (RJMCMC) [Green, 1995; Dellaportas et al., 2002] with uniform priors over models and all parameters (see Methods) to infer the posterior probability associated with each of the three possible developmental histories in Fig. 1A. This approach produces posterior distributions on each parameter and model index, describing the probability of different mechanisms given the data [Kirk et al., 2013]. We validated this modelling and inference approach with a set of synthetic observations compatible with different mechanisms of variance generation through development and between generations, including cases distinguishing the likely presence of an early germline (Supplementary Fig. S1), and confirmed that inference results were stable across different MCMC chains (Supplementary Fig. S2). Because the statistical approach considers individual heteroplasmy observations rather than losing information with summary statistics like the sample variance, substantial statistical power is retained to infer parameters and select mechanisms [Giannakis et al., 2023].

Fig. 2 shows the inferred posteriors for the number of effective segregation events at different stages of plant development and between generations, integrated over the different model structures in Fig. 1A. As above, this value is the number of binomial cell divisions that would be required to generate the observed heteroplasmy variance, given an effective population size of 50 mtDNAs or 7 ptDNAs per cell.

**Figure 2.**
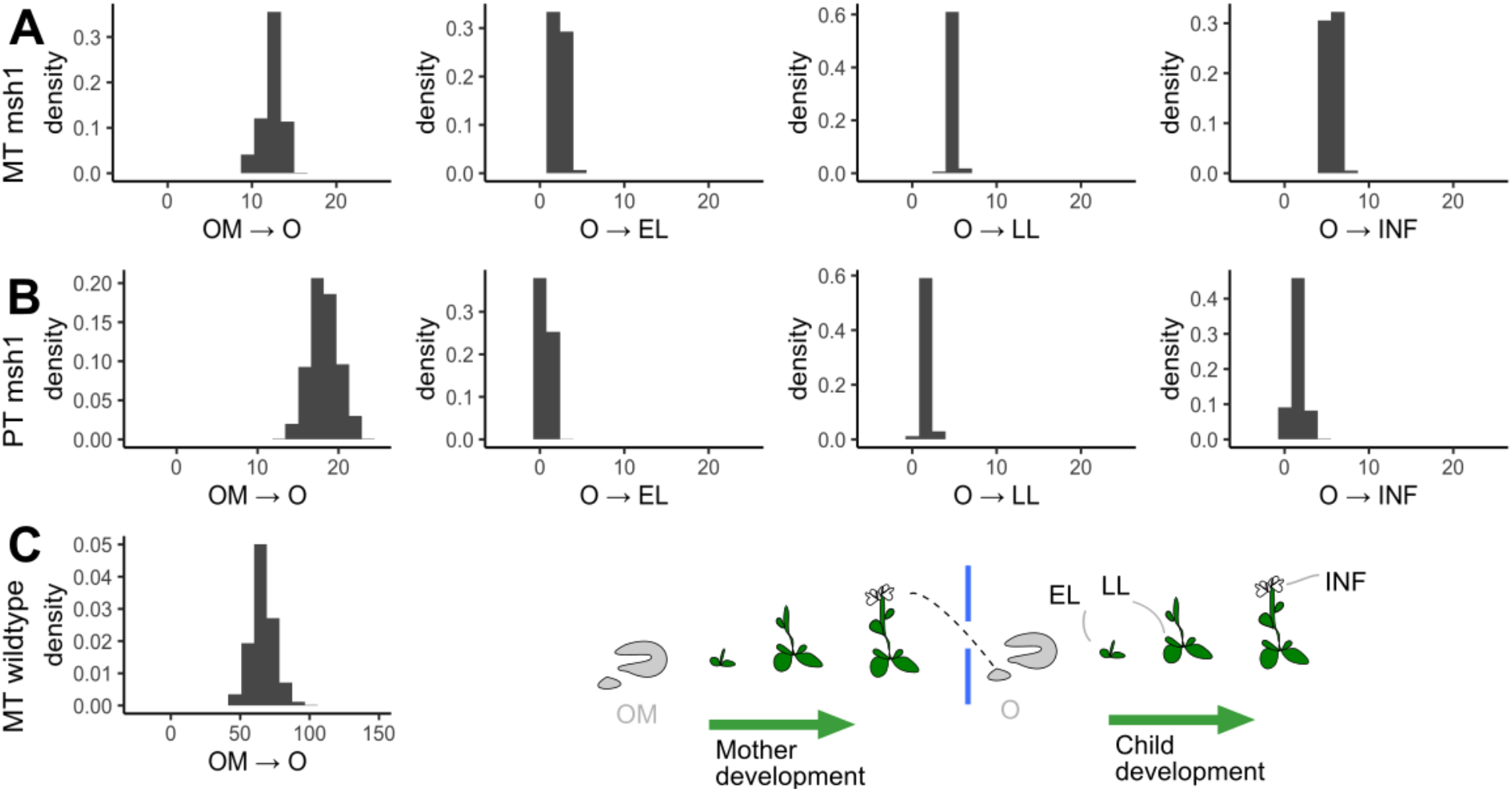
Posteriors from inference process. Posterior distributions, inferred across models, for the effective segregation events from a precursor state (O) to different tissue precursors (EL, early leaf; LL, late leaf; INF, inflorescence), and between generations (OM → O): (A) *msh1* mtDNA (*N_e_* = 50), (B) *msh1* ptDNA (*N_e_* = 7); (C) wildtype mtDNA (*N_e_* = 50, different scale).

The amount of segregation occurring between generations (OM→O) is substantially greater than that occurring within a single plant up to the inflorescence stage (O→INF). In the *msh1* mutant, a total of between 9 and 15 events are inferred to occur for mtDNA and between 15 to 25 for ptDNA between generations. In the wildtype, between 50 and 100 events – on average around a seven-fold increase in segregation -- are inferred to occur between generations for mtDNA. These numbers correspond to normalised heteroplasmy variances V’(h) of 0.17-0.26 for *msh1* mtDNA, 0.90-0.98 for *msh1* ptDNA, and 0.64-0.87 for wildtype mtDNA; where the usual “bottleneck size” is 1/V’(h). In all cases, substantial segregation is inferred to occur between the bulk inflorescences of one generation and the early stem cells in the next. This could correspond to the generation of large cell-to-cell variability within the reproductive cells in an inflorescence, matching the generation of variance in female reproductive cells in mammalian systems.

Segregation differences in samples within a generation were less pronounced, with comparatively few variance-generating events inferred to occur up to the generation of early leaves (sampled at 4-5 weeks of growth), and few more inferred to occur up to late leaf generation (sampled at 8 weeks of growth). The means of each posterior show a roughly linear trend through within-plant development, with heteroplasmy variance increasing through developmental stages; but the extent of this increase is at most half the total segregation between generations.

Due to sampling limitations in Broz et al. [2022], no within-plant samples were generated for wildtype mtDNA, and *msh1* ptDNA sampling was also somewhat limited. Based on the seven-fold scaling of mtDNA segregation from the *msh1* mutant to the wildtype, we hypothesised that the amount of segregation at each within-plant developmental stage would also be scaled seven-fold. We next set out to test this prediction and to verify the results of the ptDNA inference with further experiments.

### New heteroplasmy observations support and refine model predictions for segregation dynamics

To further illuminate the developmental dynamics of *Arabidopsis* heteroplasmy, we measured mitochondrial heteroplasmy across developmental profiles in lines where MSH1 functionality was recovered by back crossing to a wildtype male, while preserving the heteroplasmy that was present in the female. The heteroplasmy dynamics in these lines are expected to reflect those in the wild type (where heteroplasmy rarely arises because of low mutation rates and the rapid sorting). The new observations are shown in Fig. 3A-B.

**Figure 3.**
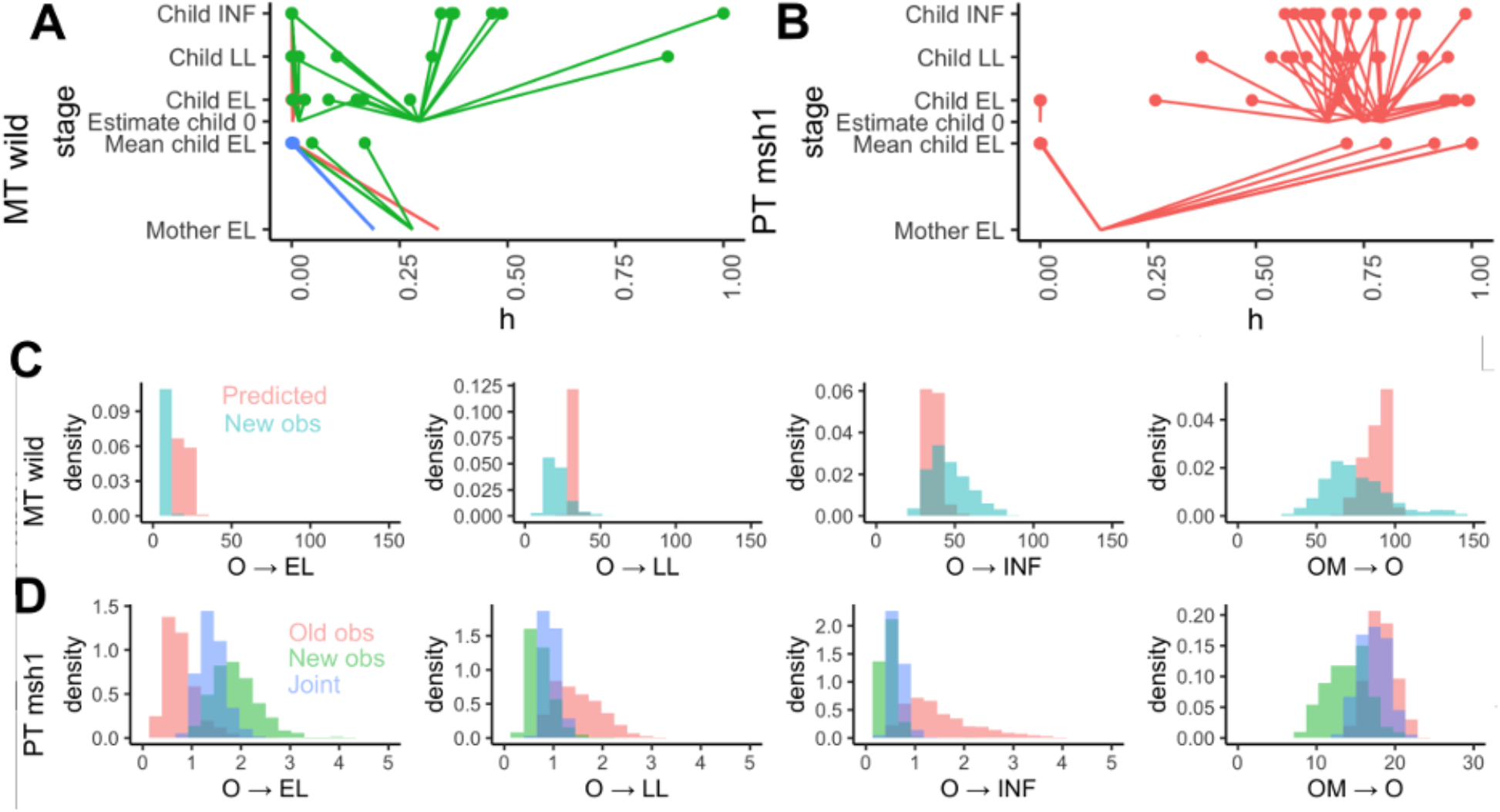
New data and predicted segregation behaviour. (A-B) New oDNA observations for (A) wildtype mtDNA and (B) *msh1* ptDNA, displayed as in Fig. 1C-E. (C) Within-plant segregation dynamics for wildtype mtDNA. Predictions (red) from scaling the *msh1* observations seven-fold to match between-generation observations; (blue) inferred effective segregation events from new data. (D) Segregation dynamics of *msh1* ptDNA; previous observations (red); new observations (green); and refined posteriors inferred from the joint dataset (blue).

Matching our predictions, we found dramatically accelerated mtDNA segregation in the wildtype at the late leaf and inflorescence stages, with the rates inferred from new observations compatible with the seven-fold scaling predicted from the between-generations data (Fig. 3C). However, the extent of wildtype mtDNA segregation prior to early leaf development was lower than this hypothesis predicted – and more similar to the lower levels in the *msh1* mutant. This difference suggests a refinement to our predicted picture -- that the increased segregation activity of MSH1 is mainly manifest in later development, which in turn is in qualitative agreement with observed patterns of MSH1 expression (Supplementary Fig. S3).

Our new ptDNA observations also matched the predictions inferred from previous data, with the increased volume of observations substantially refining the estimates of variance-generating events at different developmental stages (Fig. 3D). The new observations were always compatible with the (more uncertain) inferred posteriors from the original measurements, and combined provide a tightly defined estimate of segregation dynamics through development. Assuming as before an effective population size *N_e_*= 7, the number of variance-generating events is quite limited from early leaf to late leaf to inflorescence, with an over ten-fold further increase in segregation following between generations. It seems likely that this dramatic segregation between generations is due to a severe physical bottleneck on ptDNA, perhaps involving the inheritance of only approximately one homoplasmic organelle (see Discussion).

To ask whether within-generation segregation was a genuinely continuous process, we next explored the probability that the magnitude of segregation increased sequentially through developmental stages (for example, whether the amount of segregation experienced by late leaves exceeded that experienced by early leaves). Here we found evidence for continuous segregation through development when functional MSH1 is present, but more limited support when MSH1 was compromised (Fig. 4A). When MSH1 is compromised, segregation patterns can be explained by all segregation occurring in early development prior to early leaf sampling; with functional MSH1, segregation proceeds continuously through development.

**Figure 4.**
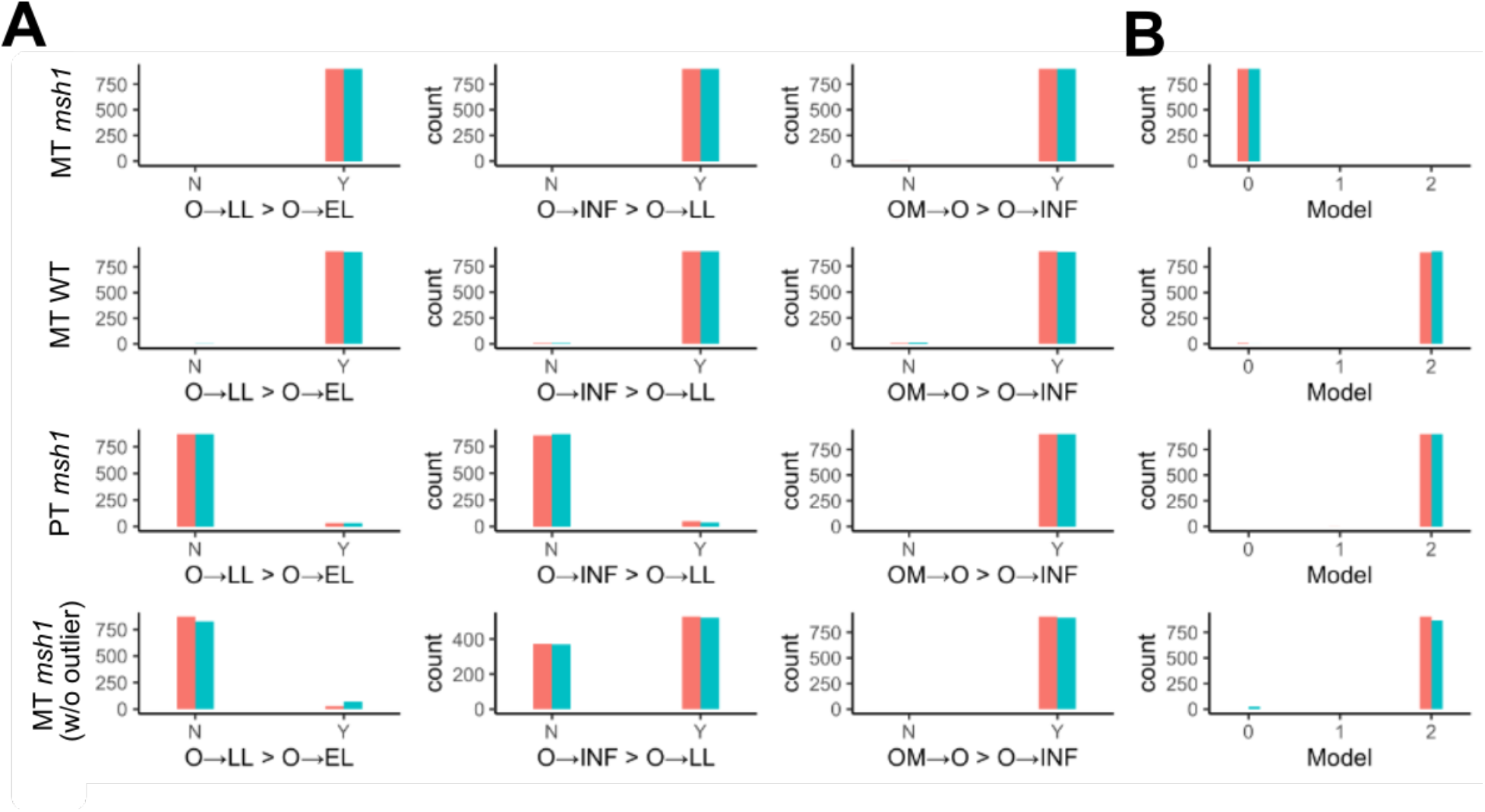
Patterns and models of segregation through development inferred from combined heteroplasmy profiles. (A) Evidence for progressive segregation through development. Posterior support for the magnitude of segregation at different developmental stages increasing sequentially – for example, O->LL > O->EL asks whether late leaves have experienced greater segregation than early leaves. (B) Posteriors on model index from reversible jump MCMC. Models 0-2 are respectively the linear germline, separate soma, all separate lineage models from Fig. 1. Rows correspond to different organelle-mutation combinations: ; the final row is the mtDNA *msh1* mutant with one potentially outlier lineage removed (see text). The two colours correspond to results from different RJMCMC simulation to demonstrate convergence (see also Supplementary Fig. S2).

### Cell divisions account for oDNA variance in the *msh1* mutant, and gene conversion can account for additional wildtype segregation of mtDNA

*Arabidopsis* has been estimated to undergo around 34 germline cell divisions between generations [Watson et al., 2016]. In the *msh1* mutant, the number of inferred effective segregation events (averages around 12 for mtDNA and 20 for ptDNA) easily fall within what would be expected from this number of binomial cell divisions for cellular populations of *N_e_*= 50 mtDNAs and *N_e_* = 7 ptDNAs, meaning that the observed heteroplasmy variance could then be readily accounted for through random cell divisions and reamplification alone.

In the wildtype mtDNA, much more segregation is observed than can be accounted for by 34 cell divisions – the average number of inferred events is around 75. Several possibilities exist for the mechanism generating this additional variance. As hypothesised in mammalian systems, partitioning of oDNA clusters, increased random turnover of oDNA, and oDNA replication restricted to a subset of the cellular population can all increase heteroplasmy variance (reviewed in Johnston [2019b]).

However, given the clear difference between the wildtype and *msh1* mutant, we suggest that an MSH1-dependent process may be responsible for this increased segregation in *Arabidopsis*. Following Edwards et al. [2021], we propose that gene conversion may be this process – in the Discussion we consider alternative mechanisms. That reference characterised the contribution of gene conversion to V’(h) as 2(1-*f*) *κ t*, where *t* is time, *f* is the proportion of mtDNA molecules in a fused state and thus physically capable of recombination, and *κ* is the rate of gene conversion between a pair of fused molecules per unit time. As the difference between V’(h) in *msh1* and wildtype mtDNA is roughly 0.5, this expression suggests that a rate of *κ* = 0.007 per cell division (corresponding to ∼0.1 gene conversion events per mtDNA per cell division; see Methods) would be sufficient to generate the observed segregation patterns over ∼34 cell divisions.

This approach employed a linear noise approximation that may be challenged by the substantial segregation magnitudes involved in this system. To check these results, we constructed a stochastic model for oDNA during development, including binomial cell divisions, random reamplification between divisions, and a variable rate of gene conversion in a population of *N_e_*= 50 oDNA molecules (see Methods). We asked what rates of gene conversion were required to generate the observed V’(h) within ∼34 cell divisions, finding support for a figure around 0.25 events per mtDNA per cell cycle (Supplementary Fig. S4). This combined model provides predictions for heteroplasmy distributions at any given stage of plant development (Supplementary Fig. S5). We should note that this gene conversion activity could be partitioned into more intense bursts in reduced developmental stages to achieve the same variance generation – as suggested by the new mtDNA observations in Fig. 3, where early meristem development appears not to generate as much segregation as later developmental stages. Such a partition of activity would agree with observed patterns of *MSH1* expression during plant development (Supplementary Fig. S3) and the observed physical behaviour of mitochondria, forming a reticulated network in the shoot apical meristem, with the potential to facilitate recombination between mtDNA molecules [Seguí-Simarro & Staehelin, 2009; Edwards et al., 2021].

### Plant germline history

The posterior distributions we have presented are integrated over all the model structures in Fig. 1A, so that they reflect “universal” behaviour regardless of the support for the individual models. However, the RJMCMC process also quantifies this support for the different models of the plant germline. Interestingly, we initially observed some diversity in the posterior distributions over this model index (Fig. 4B). The mtDNA *msh1* data has strong support for the “linear germline” model, while the mtDNA wildtype and ptDNA *msh1* data provide strong support for the “all separate lineages” model (Supplementary Fig. S2).

To interpret these findings, it helps to consider the behaviour of heteroplasmy statistics under the different models. Under separate lineages, samples from different developmental stage (EL, LL, INF) reflect the mean heteroplasmy of an early stem cell (CS1) and have independent variances. Under a linear germline, progressive sampling events form the precursor state for each stage. Differences in heteroplasmy mean can therefore arise due to this sampling, and the variance for each stage is “overlaid” on top of any such mean variability (Fig. 1B). Observations where mean heteroplasmy shifts between developmental stages are therefore more compatible with a linear germline model; limited or no shifts in mean heteroplasmy may select the separate lineages model as more flexible.

The mtDNA *msh1* data has one developmental lineage in particular that suggests a strong shift in mean heteroplasmy (top right of Fig. 1C), where an initial heteroplasmy around 0.85 gives rise to several homoplasmic late leaves and inflorescences. If this lineage is removed from the dataset, the results of inference fall more in line with the other systems (Fig. 4). If that lineage is regarded as an outlier corresponding to an accident of sampling – where, for example, other heteroplasmic inflorescences may have existed to bring the mean heteroplasmy back down – then all the remaining data support a model where segregation proceeds independently in different tissue types after diverging from a developmentally early source. There is thus at least some support for the heteroplasmy profiles in inflorescences and leaf tissue developing independently [Lanfear, 2018], although further characterisation of somatic heteroplasmy in wildtype lineages will help resolve this question.

## Discussion

We have shown, with a combination of oDNA measurements from heteroplasmic plant lines and mathematical modelling, how oDNA segregation proceeds through plant development and between generations (Fig. 5). New experiments support the predictions of the inferred mathematical models; the models make further predictions about heteroplasmy distributions at any stage of plant development (Supplementary Fig. S5). We have shown that in the absence of MSH1 functionality, oDNA segregation can largely be accounted for by the physical process of binomial partitioning at cell divisions. Although other mechanisms likely support some gene conversion activity in the absence of MSH1, high rates of such activity are not required to explain observed segregation patterns in the mutant. By contrast, MSH1 functionality induces a seven-to ten-fold increase in segregation strength, leading to rapid shifts towards homoplasmy, which cannot be explained by cell divisions alone.

**Figure 5.**
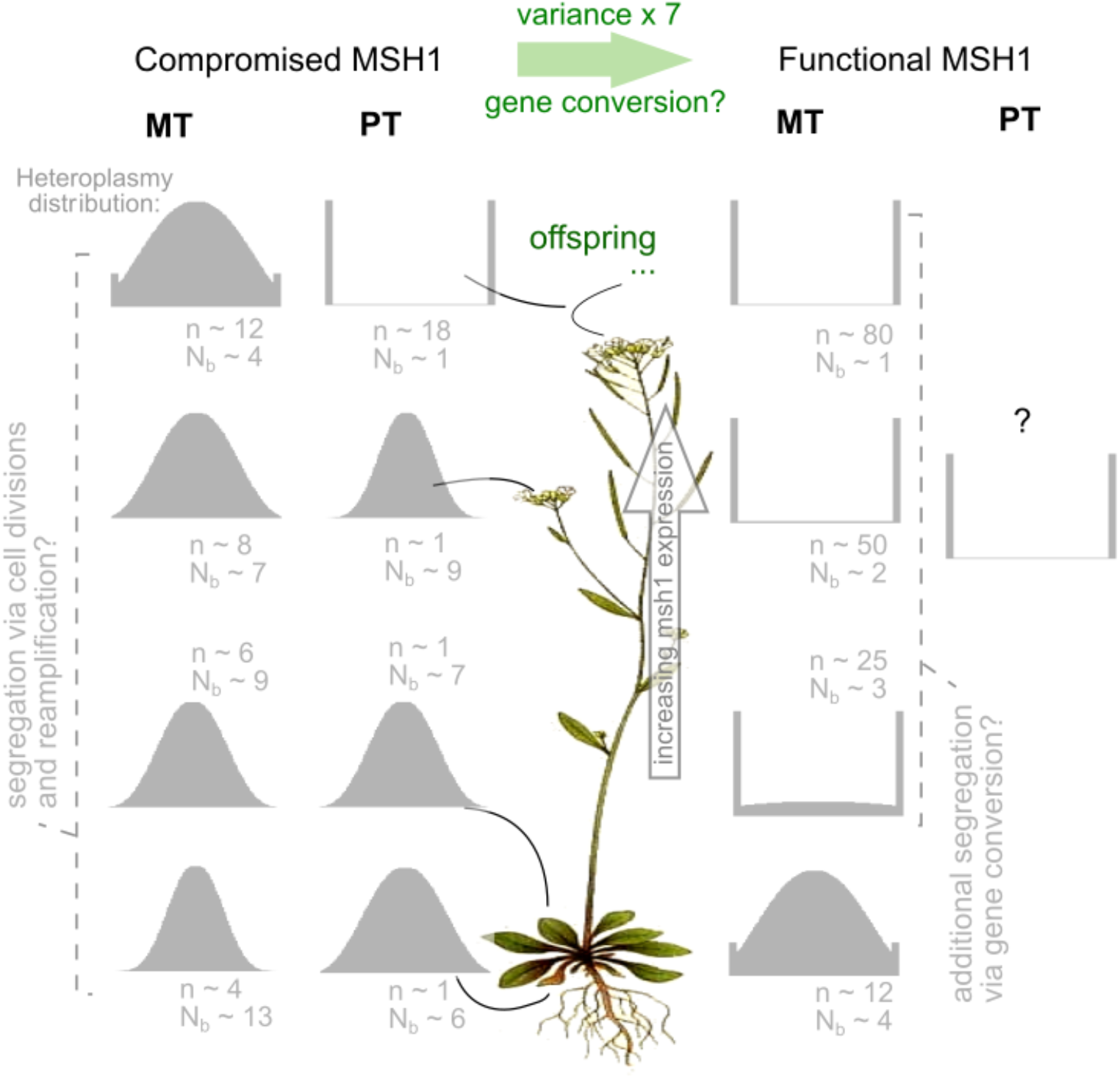
Summary of inferred segregation dynamics within plants and between generations. Illustrative distributions of heteroplasmy, corresponding to the inferred mean segregation magnitude (*n* segregating events, for *Ne* = 50 mtDNAs or *N_e_* = 7 ptDNAs; and *N_b_*, effective bottleneck size). Distributions at each developmental stage, and an initial heteroplasmy of 0.5, are shown for mtDNA (MT) and ptDNA (PT) in wildtype and *msh1* mutants (all wildtype PT observations are homoplasmic, so no inference is possible; see Discussion for hypotheses). Continuous segregation is supported by inference in all systems except PT; model selection suggests most support for a picture where separate developmental lineages are involved for each developmental stage.

We do not have measurements of heteroplasmic ptDNA on the wildtype background – all lines measured so far have been homoplasmic. The predictions of this theory for wildtype plastid heteroplasmy dynamics depend on the spatial arrangement of ptDNA information. If ptDNA within a single plastid is homoplasmic, and heteroplasmy arises from a mixture of internally homoplasmic organelles, then the effect of functional gene conversion will be limited. This is because each ptDNA will usually only be physically colocalised with an identical partner, leaving no capacity to change genetic identity. If, however, plastids are internally heteroplasmic, functional gene conversion may act to further speed up segregation. In this case, following observations for mtDNA, we would expect roughly seven times as many effective cell divisions to take place (matching the mtDNA case), leading to an effective 150-200 cell divisions for the *N_e_* = 7 case. This would lead to homoplasmy in all but a very small proportion of offspring (as observed).

The quantitative details of our model depend on some assumptions, including a binomial division – random reamplification model for oDNA at cell divisions, the Kimura model for oDNA heteroplasmy, and particular choices for effective population size of oDNAs. The choices we have made have support from the literature (see Methods), but are not expected to be universally true or perfectly precise single values. oDNA population sizes change through development (see Methods and references therein) and oDNA partitioning at cell divisions may be more or less tightly controlled than a binomial distribution [Jajoo et al., 2016; Johnston et al., 2015]. Our effective ptDNA population size is based on a picture where ptDNA populations inside individual plastids are homogeneous: this assumption may be challenged in the case of recent *de novo* mutations that have not yet fixed within an organelle. The results we report – the relative magnitudes of segregation at different developmental stages, the difference between wildtype and *msh1* lines, the role for gene conversion, and the agreement of new experiments with theoretical predictions – are robust with respect to different choices of these parameters. The specific numbers of segregating events we infer should be interpreted as effective quantities, reflecting biological reality if our parameter choices are accurate, otherwise requiring some scaling (see Methods and Supplementary Fig. S6) for a precise quantitative connection to other conditions.

The indirect evidence from our study, after removing a potential outlier lineage, supports a picture where different tissues, including the germline, have different developmental lineages after diverging from a developmental ancestor [Lanfear, 2018]. A previous study in carrot [Mandel et al. 2020] did not find shifts of mean heteroplasmy during development, although some individual observations suggested the capacity for large minor allele amplification; this picture would also be compatible with segregation (increasing variance) through independent developmental lineages. Regardless of the within-plant model, most of the between-generation segregation we observe occurs between the inflorescences of the mother and the early meristem of the offspring. For plastids in particular, it seems likely that this strong segregation may be in part due to a physical bottleneck, where a small number – perhaps just one in some cases – of homoplasmic organelles are inherited. For mitochondria, our observations support a picture where some segregation occurs progressively through development. The rate of this increase is limited in the *msh1* mutant and clearer in the wildtype, and its magnitude is smaller than the between-generation shift.

Substoichiometric shifting (SSS) involves the sudden amplification of a rare mtDNA type (a sublimon) to dominance [Abdelnoor et al., 2003; Arrieta-Montiel et al., 2001; Woloszynska, 2010]. The dynamics characterised here illustrate how this amplification may occur. Even if a sublimon is present only rarely in SAM cells, if one of those cells becomes the precursor to a plant branch or organ, the sublimon can very naturally (and quickly) come to dominate that branch or organ (and hence offspring from it). Our work here quantifies how this shifting may occur across different organs in a plant, leading to inherited differences. In a similar vein, branch-to-branch differences in variegation caused by oDNA features have been recognised for over a century (initially laying the foundation for the understanding of cytoplasmic inheritance [Hagemann, 2010]). Such branch-to-branch differences are caused by the segregation of oDNA from an initially heteroplasmic state across different parts of the plant. The quantitative model we present links, for example, the unobservable initial inherited heteroplasmy to the proportion of different variegated phenotypes throughout the plant, by quantifying the extent of segregation through different periods of plant development.

Observations here and in Broz et al. [2022] point to MSH1 dramatically accelerating oDNA segregation. We have proposed that this acceleration may be due to gene conversion. However, the function and mechanism of action of MSH1 in plants remain debated. Evidence certainly points to its role in the control of oDNA recombination (often described as recombination surveillance [Abdelnoor et al., 2003; Shedge et al., 2007]). Its unusual structure - - including an endonuclease domain - - has led to the suggestion that it induces double stand breaks that then provide the substrates for gene conversion [Christensen, 2014]. The heteroplasmy measurements here strongly suggest that MSH1 acts to generate high cell-to-cell variance in oDNA heteroplasmy through plant development. Theory has suggested gene conversion as one plausible mechanism with desirable properties [Edwards et al., 2021]. However, it may be that MSH1 generates heteroplasmy variance via another mechanism. Depletion of oDNA copy number, for example, would impose a physical bottleneck on the population, both amplifying variability from divisions and inducing variability from subsampling the population. If MSH1 acts to deplete oDNA, these effects could be of comparable or greater importance in generating variability, depending on the quantities involved [Cree et al., 2008; Johnston et al., 2015]. Broz et al. [2022] showed that oDNA copy number was not significantly impacted in leaves of MSH1 versus wildtype plants, but it is unknown whether these results reflect oDNA levels in germline. If, in some way, MSH1 enforces replication of a subset of oDNA molecules as proposed by Wai et al. [2008] in a mammalian context, this mechanism could also explain the observed segregation. While the evidence points towards a more direct link between MSH1 and gene conversion [Wu et al., 2020; Broz et al., 2022], we cannot completely discard these hypotheses without measurements of copy number and oDNA replication activity. We were unable to find or acquire estimates for absolute rates of oDNA recombination in *Arabidopsis*; future estimates of these quantities will help provide further evidence for these mechanisms. It is noteworthy that *MSH1* expression is increased relative to other tissues in the meristem in *Arabidopsis* and other species (Supplementary Fig. S3, [Edwards et al., 2021]), and that mitochondria physically fuse to a greater extent in the meristem cells [Seguí-Simarro & Staehelin, 2009; Edwards et al., 2021]. Physical colocalization of mitochondria is a prerequisite for mtDNA interaction and recombination [Logan, 2006; Arimura, 2018; Giannakis et al., 2022], and the collective dynamics of mitochondria are altered in the *msh1* mutant, potentially as a compensatory response to support more interaction [Chustecki et al., 2022; Chustecki et al., 2021]. Together, these observations suggest a linked physical and genetic axis of control acting to shape oDNA through plant generations.

## Methods

### Plant material and growth

The initial generation and selection of heteroplasmic plant lines is described in Broz et al. [2022]. Here, plants of the homozygous *msh1* (At3g24320) mutant line CS3372 (*chm1-2*) were used for analysis of plastid heteroplasmy. For mitochondrial heteroplasmy analysis in a wild type background, maternal lines of *msh1* CS3246 (*chm1-1*) were crossed with wildtype males to generate F1 progeny. These different *msh1* alleles were used because it was on these backgrounds that oDNA variants present at reasonable allele frequencies arose and were retained (see below); both have been reported to be full allelic knockouts [Broz et al., 2022]. All progeny were confirmed to be heterozygous for MSH1. Seeds of desired lines were vernalized in water at 4 °C for 3 days, sown in 3 inch pots containing Pro-Mix BX media and grown under short day conditions (10 h light / 14 h dark) on light racks with fluorescent bulbs (∼150 µE m^-2^ s^-1^) at ambient temperature (∼25 °C). An initial fully expanded rosette leaf sample was taken at 4 weeks of growth to identify heteroplasmic individuals. Three additional leaves were sampled at 5 weeks of growth. These 4-5 week old leaf samples are considered “early leaf” (EL) for subsequent analyses. At 8 weeks, four additional leaf samples were taken. Two were harvested from the base of the rosette. These leaves were already fully expanded at 5 weeks and emerged from the SAM around the same time as the EL samples described. Thus, these are also considered “EL”. Two additional fully expanded leaves were harvested at 8 weeks from the top of the rosette, emerging from the SAM at a later timepoint that ELs, and are considered as late leaf “LL” in the analysis. Inflorescence tissue (INF) was harvested after plants began to bolt.

### Heteroplasmy measurements

DNA extraction and heteroplasmy analysis were performed as described previously [Broz et al. 2022]. Briefly, single nucleotide variants (SNVs) in oDNA of *msh1* mutant lines were identified by sequencing [Wu et al. 2021] and ddPCR assays were designed to track these SNVs within plants and between generations. Allele specific primers and probes were designed to each SNV and droplet generation and reading was performed using Bio-Rad QX200 system. This study used the specific loci plastid 26553, mitochondria 91017 and mitochondria 334038, which were retained after screening the original set of heteroplasmic variants for those present at moderate allele frequencies. A correction factor was applied to mitochondrial data to account for the amplification of nuclear copies of the mitochondrial genome (numts) found in Arabidopsis. Specifically, the large numt on chromosome 2 is too similar to actual mtDNA to be distinguished with short reads or ddPCR markers. So we approximate the number of nuclear genome copies in the sample (which would inflate the number of apparent mitochondrial “wild type” alleles) and correct accordingly. All nupts have enough sequence divergence that nuclear and plastid copies can be unambiguously distinguished.

### Developmental history models

First picture a fertilised zygote giving rise to an early population of stem cells. At some developmental time point this population will contain the single ancestral cell of all early leaf samples, as well as of cells that will continue to proliferate in the SAM. At a later time point, the new SAM population will contain the ancestor for all late leaf samples, as well as for further proliferating cells. At a still later time point, the new SAM population will contain the ancestral cell to all inflorescence samples. Inflorescences are interpreted as containing the egg cells for the next generation, in which the developmental outline above is repeated for each single fertilised zygote. Each tissue’s heteroplasmy value is drawn from a distribution describing some amount of segregation acting on developing descendants of these ancestral stem cells, with relationships described via the “cell pedigrees” or “lineage trees” in Fig. 1A [Wilton et al., 2018; Stadler et al., 2021].

The developmental history of plant germlines is debated [Lanfear, 2018]. To compare hypotheses on plant germline behaviour, we also consider two additional alternative models. In Fig. 1B, the future germline is sequestered early in development and then develops in parallel to the somatic tissues. Here, the model is as above, except the inflorescence ancestral cell is drawn from the early stem cell population. In Fig. 1C, separate somatic lines also exist, so that the different organs all develop independently from an original early precursor. In theory, different germline histories – where soma and germline are sequestered at different developmental timepoints – will give rise to different correlations and variance structures in the oDNA populations in different tissue types. For example, if the germline develops independently of the soma, correlations between mean oDNA heteroplasmy in somatic and inflorescence samples are less likely, and it may be possible for inflorescence oDNA to have lower variance than soma oDNA. If the germline shares a common developmental ancestry with the soma, correlations are more likely, and inflorescence variance will be at least as high as soma variance.

### Inference of segregation dynamics

To assign a likelihood to our tissue observations given a developmental model, we need to (a) estimate the ancestral cell heteroplasmies and (b) estimate the probability of observing a tissue heteroplasmy given the ancestral value and some parameterised description of segregation [Burgstaller et al.,2014; Burgstaller et al., 2018]. For (a), we treat ancestral cell heteroplasmies as latent variables and integrate the likelihood over all possible values for each. For (b), we use the Kimura distribution [Wonnapinij et al., 2008; Kimura, 1955] to describe the probability of observing a given heteroplasmy in individual tissue samples, creating a stochastic model with a full likelihood function [Giannakis et al., 2022b, Broz et al., 2022]. We change variables from the “drift parameter” *b* to an effective number of variance-generating events *n* = log *b* / (1 – 1/*N_e_*) (see below) to provide a convenient, additive parameter for serial segregation events. The corresponding likelihood is then used in a reversible jump Markov chain Monte Carlo (RJMCMC) framework [Green, 1995; Dellaportas et al., 2002] (see below) with uninformative uniform priors on initial heteroplasmies and division numbers and compute posterior distributions over these parameters.

For numerical efficiency, we precompute Kimura distributions for 0 to 200 cell divisions and initial heteroplasmies from 0 to 1 in steps of 0.01 and use these precomputed distributions as a lookup table in the inference process. For numerical efficiency, we set effective population size to 50. A post-hoc correction can be used to interpret the results from this setup in terms of any other population size (see below).

To account for the fact that heteroplasmy measurements may have some associated uncertainty, we implement a degree of granularity within the model. For example, a granularity of 0.01 means that heteroplasmy values are rounded to the nearest 0.01. This both allows for measurement noise and improves computational speed; we will show that our results are robust to different choices of this parameter.

We write {D_i_} = {D_i,ME_, D_i,CE_, D_i,CL_, D_i,CI_} for the set of observations in family i, with elements respectively corresponding to Mother Early leaf, Child Early leaf, Child Late leaf, and Child Inflorescence. We write S_Cj_ for the latent variable associated with ancestral cell heteroplasmy at developmental stage j. The likelihood associated with measurements, in the model without a segregated germline, is then

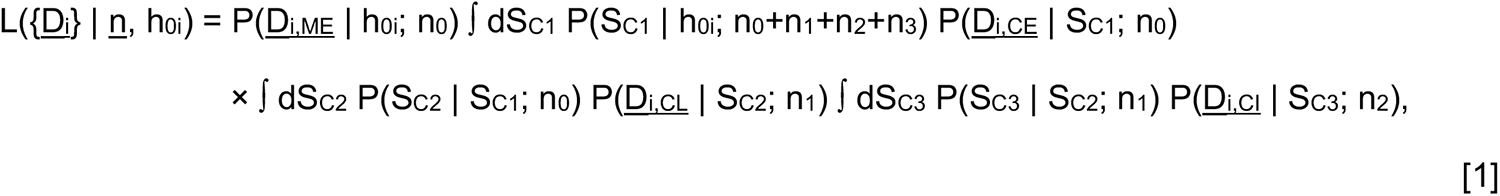

So that SC1 is the precursor to EL and SC2, SC2 is the precursor to LL and SC3, and SC3 is the precursor to INF (Fig. 1A). With a segregated germline the corresponding expression is

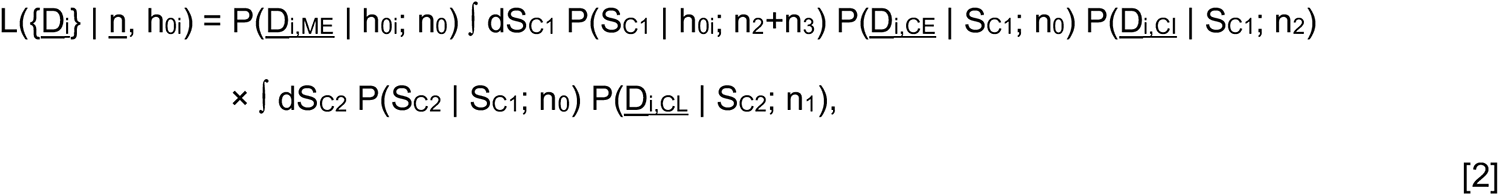

So that SC1 is the precursor to EL, INF, and SC2, and SC2 is the precursor to LL. With completely separate developmental lineages we have

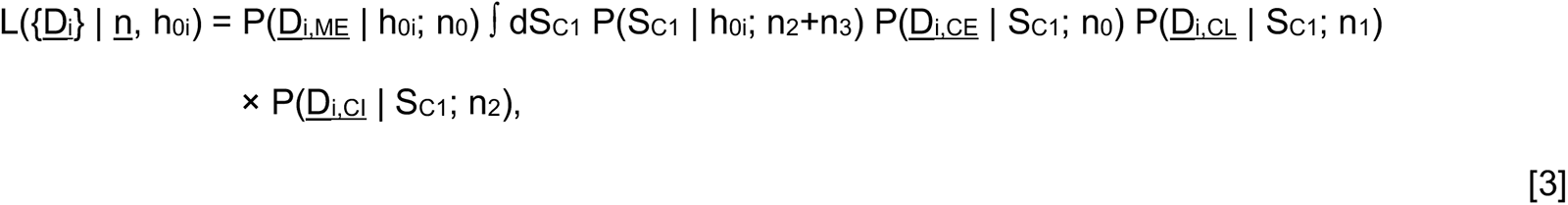

So that SC1 is the precursor to all lineages, which develop independently.

An important difference between the models is whether samples at different stages can have different population means. In the separate lineages model, EL, LL, and INF pedigrees all come from the same precursor, so have the same population mean. In the linear model, each pedigree begins with a (latent) sample from a previously segregated population (Fig. 1B), so population means can differ (Supplementary Fig. S1). They also differ in the accumulated amount of segregation at the population level. The “linear germline” model enforces a monotonic increase in segregation (hence in V’(h)) through development – hence EL ≤ LL ≤ INF ≤ cross-generation. The “all separate” model supports a more flexible picture where INF < EL, for example. However, although these relationships hold statistically at the population level, a given set of samples may not reflect them: for example, a sample of inflorescences may not capture the full possible spread of values and may thus suggest a lower variance than the true case. The full likelihood-based inference process below accounts for these sampling issues.

Given one of the above likelihood functions for a family set of observations {D_i_}, the likelihood associated with a full set of observations is

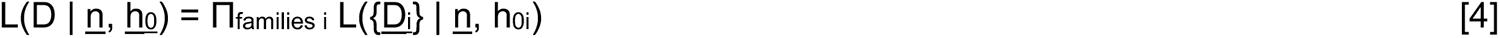

### Effective population sizes

Preuten et al. [2010] find 50 or fewer mtDNAs in stems and flowers. Wang et al. [2010] found egg cells from Arabidopsis to possess 59.0 copies of mtDNA on average. Gao et al. [2018] do not quantify mtDNA molecules in *Arabidopsis* but observe around 250 mtDNA nucleoids in mature eggs and mature zygotes, and 100-200 mtDNA nucleoids per cell during embryogenesis, with a doubling between early apical cells and mature apical cells. We choose an effective population size of 50 for consistency with those studies where mtDNA copy number is more directly observed.

In a comprehensive survey across species, Greiner et al. [2020] report an increase in plastids per cell in *Arabidopsis* development from 4-10 in the meristematic region, through 22-34 in young leaves, to 50-90+ in mature leaves. Corresponding ptDNA counts per plastid (per cell) are given as 8-21 (71-146), 48-84 (997-2476), 79-121 (2900-5500+). We choose an effective population size of 7, corresponding to the central estimate for the meristematic observations, and assuming that plastids are internally genetically homogeneous [Scarcelli et al., 2016]. This assumption may be challenged in the case of recent mutations (see Discussion).

For numerical convenience we used a population size of *N_e_*= 50 in the numerical simulations. Following the usual parameterisation of the Kimura distribution for mtDNA work [Wonnapinij et al. 2010, Giannakis et al. 2023],

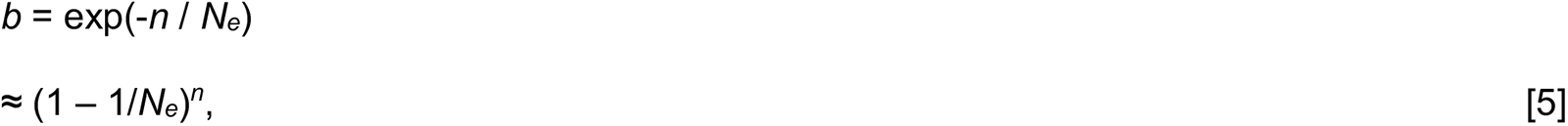

Using this approximation (which is not perfect for low *N_e_*) we can immediately interpret an inferred value of *n* for *N_e_*as equivalent to a value *n’* for *N’_e_* :

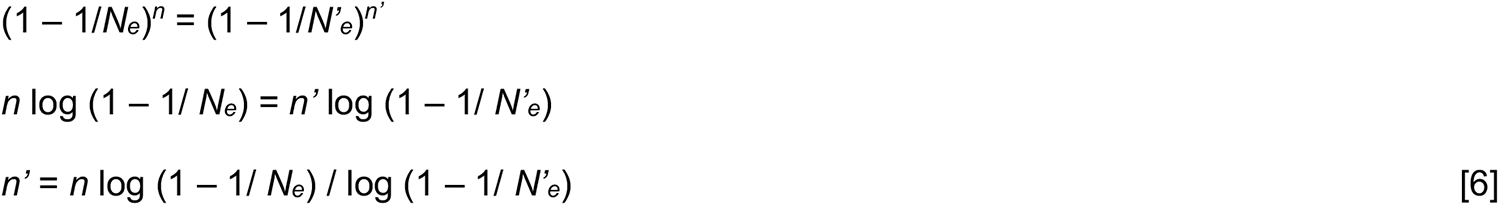

so that, for example, *n* = 10 divisions for *N_e_*= 50 give roughly the same heteroplasmy distribution as *n’* = 20 divisions for *N_e_* = 100. We can then scale the results for *N_e_*= 50, chosen for numerical convenience in our simulation, to the required effective population size in our estimates of biological reality. Hence, any of the inferred numbers *n* of segregating events we report (using *N_e_*= 50 for mtDNA and *N_e_* = 7 for ptDNA) can readily be interpreted for another effective population size *N_e_’* by multiplying by the factor log (1 – 1/ *N_e_*) / log (1 – 1/ *N’_e_*), which for most values is close to *N_e_*/*N_e_’* (Supplementary Fig. S6). Finally, effective “bottleneck size” *N_b_* (the effective population size if variance is generated by a single event) can be recovered from our inferred *n* with

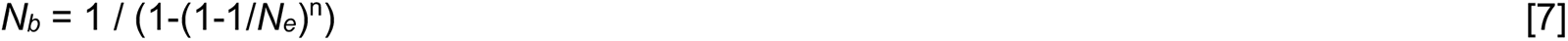

### Reversible jump MCMC

We use reversible jump MCMC to identify the support for different models of developmental histories [Green, 1995; Dellaportas et al., 2002; Kirk et al., 2013]. We explored several options for relating parameters in each model class, which all gave convergent results in the long-term limit of the MCMC chains, but found the best mixing between model classes to be achieved simply using n_i_^(1)^ = n_i_^(2)^ = n_i_^(3)^ for all developmental stages *i* and with model classes given by superscripts (1: linear germline; 2: separate germline; 3: all separate lineages), enforcing these (and preserving h_0_ values) as deterministic proposal rules upon a proposed shift from model *i* to model *j*. These expressions immediately provide the (trivial) mapping functions g_ij_(n^(i)^) for implementing such a step from model *i* to model *j* [Green, 1995; Dellaportas et al., 2002]. All models have the same dimensionality and the Jacobean determinants associated with each of these mapping functions are all one. We employ uniform priors on all parameters and model indices, making the acceptance rule for the RJMCMC implementation equivalent to the normal Metropolis-Hastings acceptance rule when a between-model step is proposed. We propose such steps with probability 1/3, employing the above perturbation to parameters when this option is not chosen. MCMC chains were run over 10^5^ samples, discarding 10^4^ as burn-in and subsequently recording every 10^th^ sample.

### Estimating and simulating variance due to gene conversion

The parameter *κ* in the main text is the rate constant associated with the gene conversion processes WT+MU → WT+WT and WT+MU → MU+MU [Edwards et al., 2021]. In a simple picture we could assume that half our *N_e_*= 50 mtDNAs are WT and half are MU. Then the rate of gene conversion is *κ* × 25 × 25, which for *κ* = 0.007 per cell division gives ∼4 events per cell division or ∼4/50 = 0.08 events per mtDNA per cell division.

The derivation of this expression depends on a linear noise approximation, and the rates in the above argument will of course vary as segregation proceeds. To provide a more precise estimate, we implemented a simple stochastic simulation of binomial cell divisions, random re-amplification, and gene conversion in a model cellular population. We simulated these processes for various gene conversion rates and 300 cell divisions and asked what gene conversion rates were needed to generate a given normalised heteroplasmy variance V’(h) within ∼34 cell divisions (Supplementary Fig. S4).

## Data and code availability

All data and code is freely available at https://github.com/StochasticBiology/plant-segregation. The inference code is written in C; the data curation and visualisation is written in R [R Core Team, 2022], using libraries readxl [Wickham and Bryan, 2022], stringr [Wickham, 2019], ggplot2 [Wickham, 2016], and gridExtra [Auguie, 2017].

## Acknowledgements

This project has received funding from the European Research Council (ERC) under the European Union’s Horizon 2020 research and innovation programme (Grant agreement No. 805046 (EvoConBiO) to IGJ). IGJ gratefully acknowledges support from the Peder Sather Center. DBS and AKB are supported by NIH Grant R01 GM118046. The authors are grateful to Ben Williams for valuable discussion.

## Supplementary Information

**Supplementary Figure S1.**
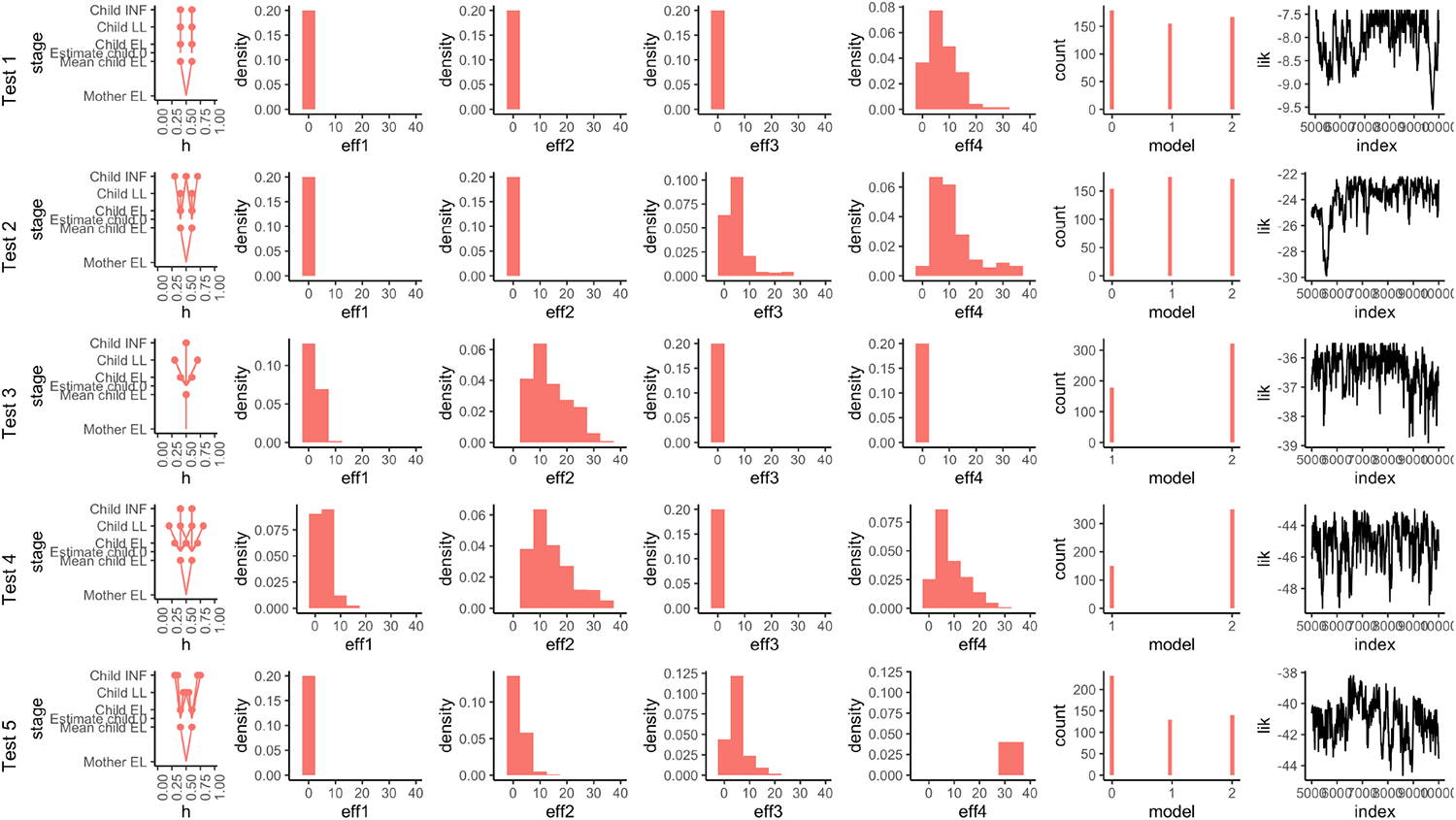
Validating model and inference approach. Each row corresponds to a synthetic dataset generated to match a different type of segregation dynamics. The synthetic observations are shown in the first column, followed by the inferred effective segregating events to EL, LL, INF, and next-generation stages (eff1-4); the inferred model index (0, linear; 1, separate germline; 2, all separate); and finally a trace of likelihood over the MCMC chain as a readout of chain dynamics. Individual experiments reflect (1) segregation between generations, generating diversity between siblings but not within plants; (2) segregation in inflorescence development (and possibly between generations) but not in somatic tissue; (3) segregation only in somatic tissue, with a separate germline; (4) segregation between generations and in somatic tissue, but with germline protected; (5) segregation throughout linear germline, with precursor cells causing shifts in mean (see Methods). In case (1), segregation between generations but nowhere else is inferred, with uniform posteriors over model index in the absence of further information. In case (2), segregation at inflorescence development but not in somatic tissue is inferred, with a linear model favoured. In case (3), zero segregation in the germline and nonzero in somatic tissue is inferred, with models 1 and 2 (separate germline) inferred. Case (4) mirrors case (3) but with between-generation segregation also inferred. Case (5) supports the linear germline model as others cannot account for the shifts in mean heteroplasmy between stages.

**Supplementary Figure S2.**
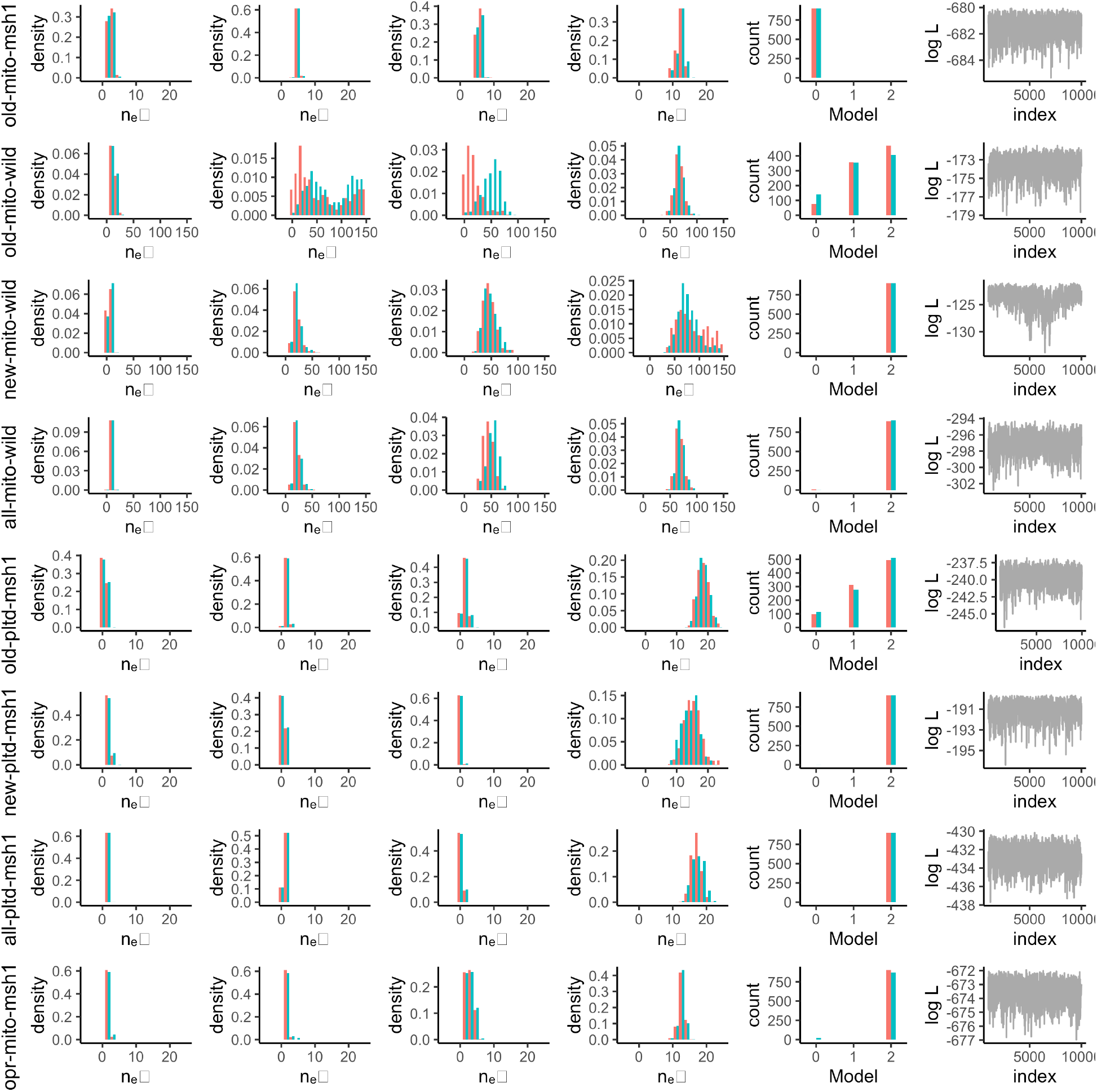
Inferred behaviour for different datasets. Each row is the result of inference on the given dataset (*old*, existing data, *new*, new data, *all*, combined data, *opr*, outlier pruned as in main text; *mito*, MT, *pltd*, PT). Effective numbers of segregating events to EL, LL, inflorescence, and between-generation stages (eff1-4); the inferred model index (0, linear; 1, separate germline; 2, all separate); and finally a trace of likelihood over the MCMC chain as a readout of chain dynamics. Results for two independent MCMC chains (red and blue) are shown for all except the likelihood traces. Divergence in the “old-mito-wild” case reflects the unidentifiability of within-plant segregation parameters from this between-generational data.

**Supplementary Figure S3.**
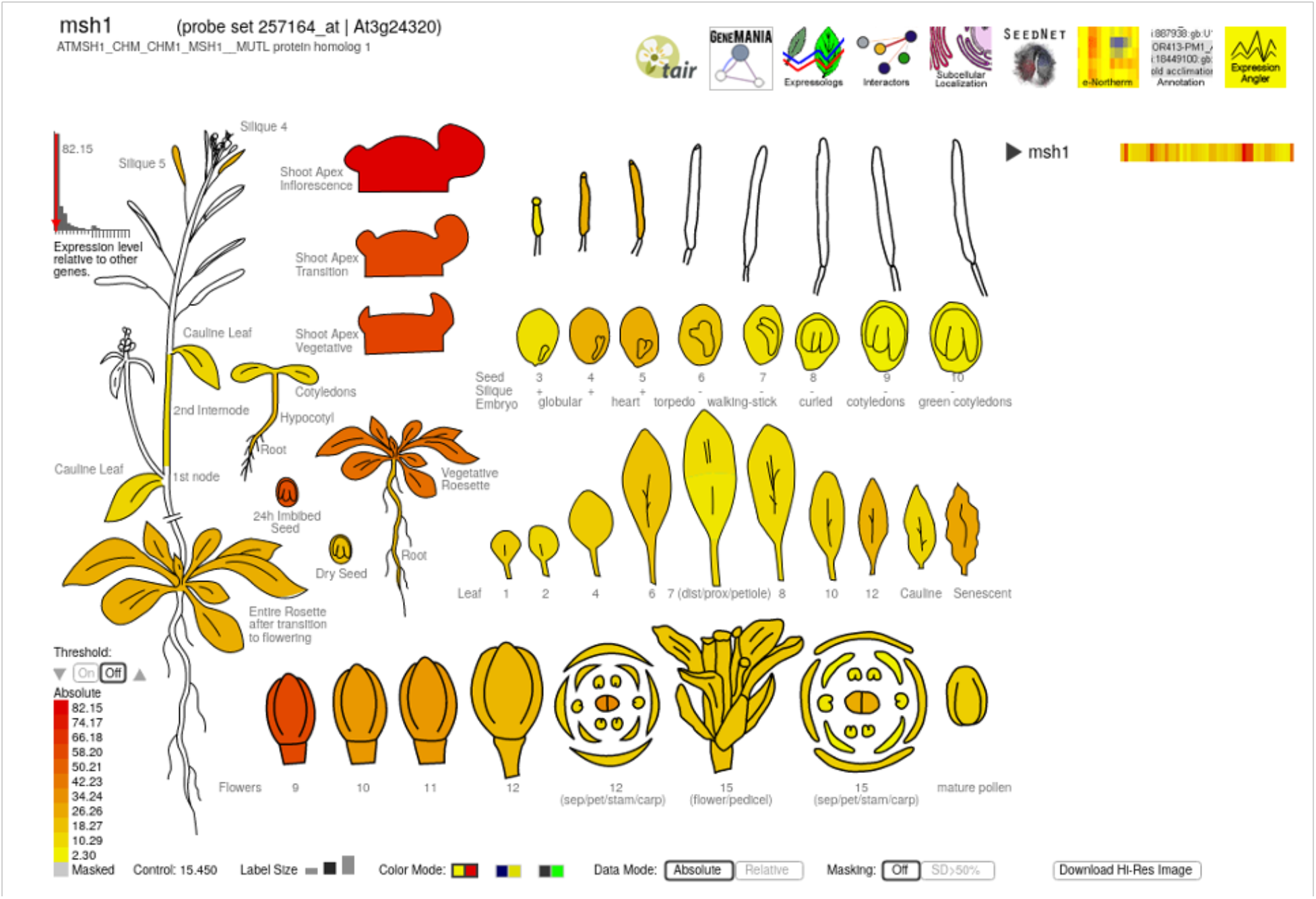
Msh1 expression patterns during development. Data from Schmid et al. [2005], visualised by the “eFP browser” from the Bio-Analytic Resource for Plant Biology [Winter et al., 2007].

**Supplementary Figure S4.**
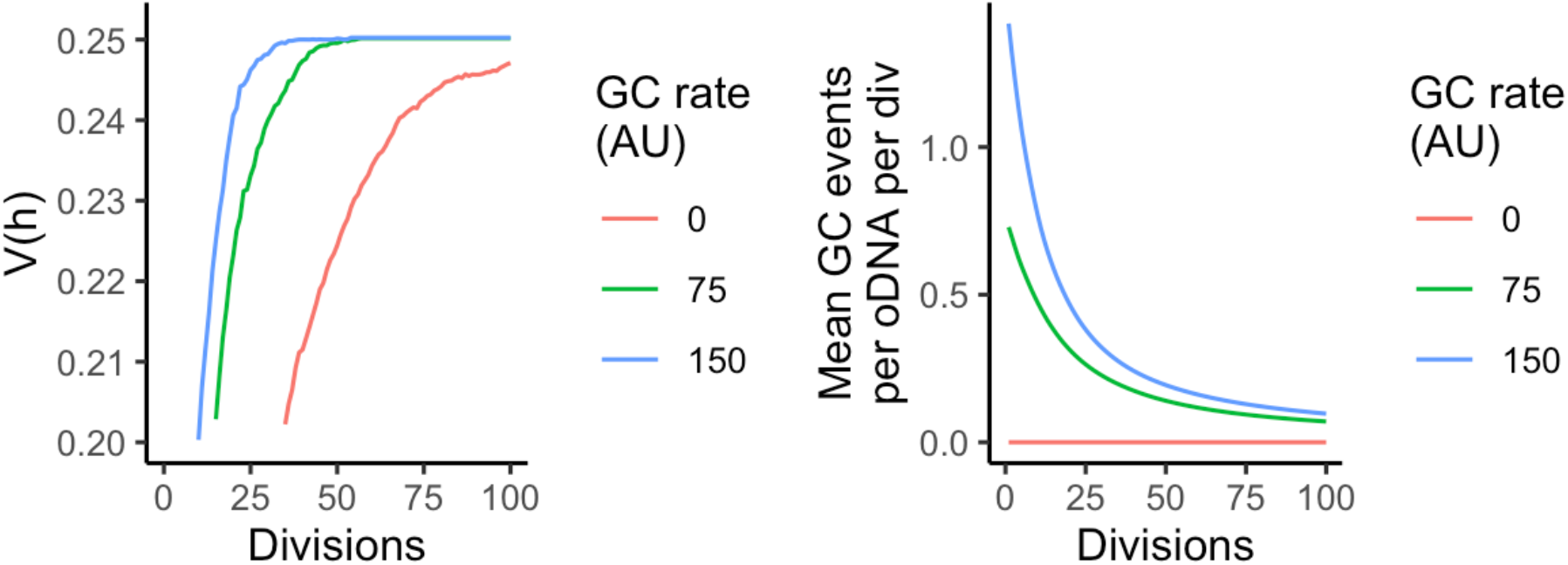
Simulated segregation with and without gene conversion. (left) V(h) with number of divisions for different rates of gene conversion attempts (GC rate). (right) Actual gene conversion events per mtDNA per division, with number of divisions for different R. Within 34 divisions, the R = 75 and R = 150 cases readily generate the V(h) ∼ 0.25 (corresponding to V’(h) ∼ 1 for these simulations where h = 0.5) values observed for 75 divisions of the R=0 case, corresponding to a mean around 0.25 gene conversion events per mtDNA per cell cycle.

**Supplementary Figure S5.**
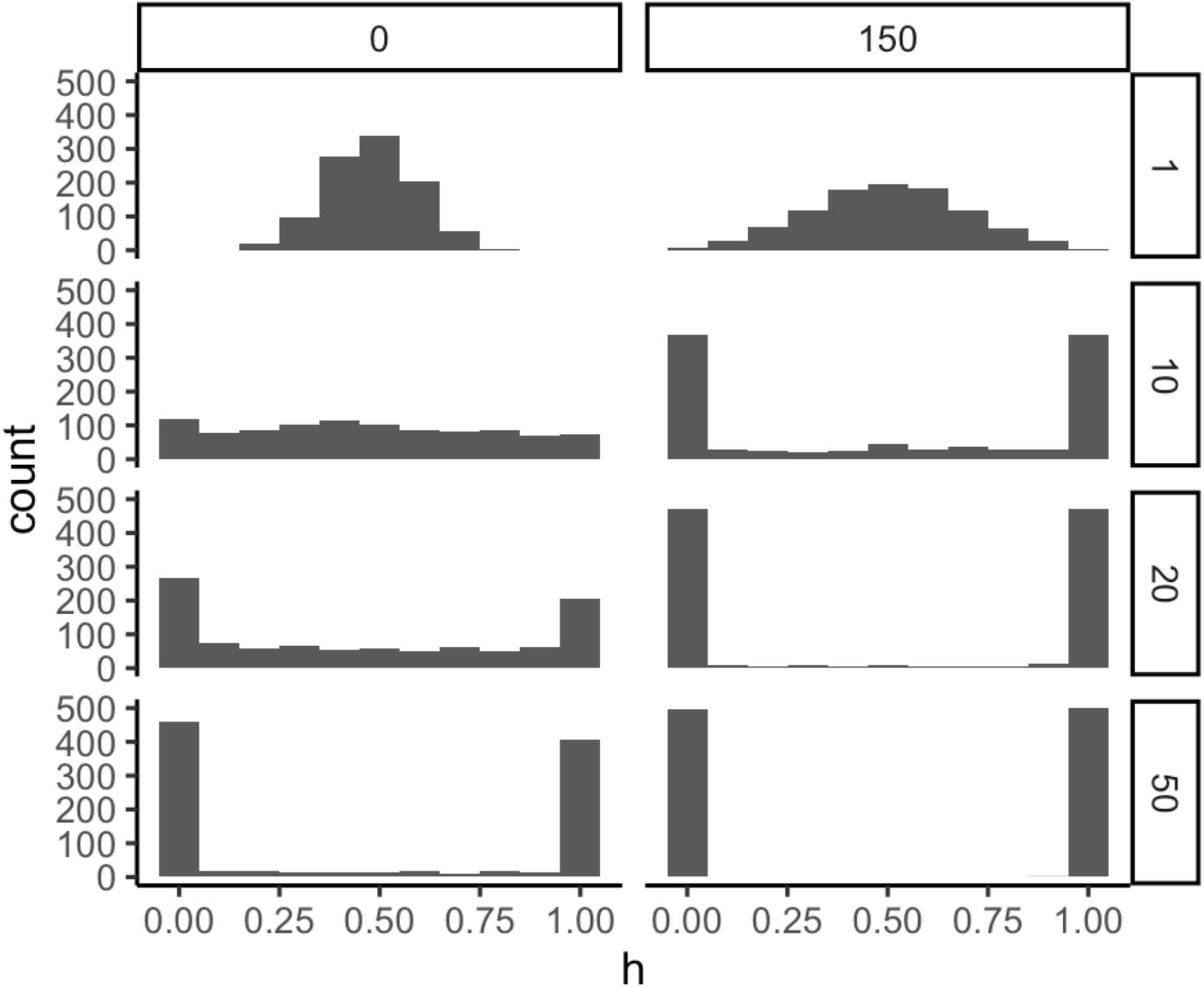
Predicted heteroplasmy distributions over cell divisions. Example model predictions for heteroplasmy distributions in mtDNA populations of size *N_e_* = 50, with a given number of cell divisions (rows). (left) No gene conversion, modelling the *msh1* case; (right) gene conversion at the rate suggested by our analysis in the wildtype plants.

**Supplementary Figure S6.**
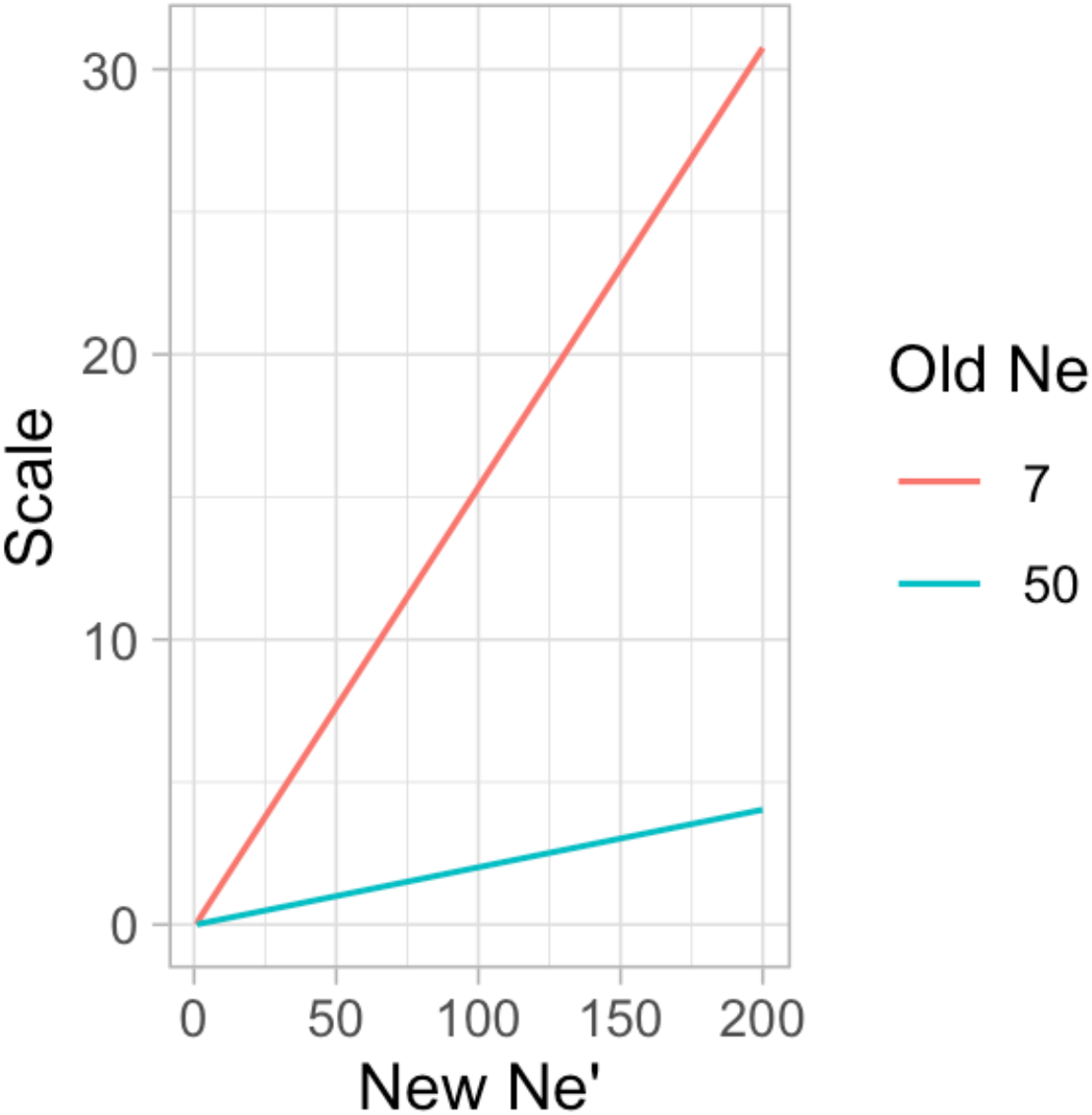
Scaling factors for converting effective population sizes. To interpret a number of inferred segregating events *n* from a population with *N_e_* = 7 or 50 with a new population size *N_e_’*, read off the scale factor corresponding to the new population size on the horizontal axis and scale *n* by this factor. For most cases this scale factor is very close to *N_e_*/*N_e_’*.

